# *Adar1* deletion causes degeneration of exocrine pancreas via Mavs-dependent interferon signaling

**DOI:** 10.1101/2021.11.03.467127

**Authors:** Dhwani N. Rupani, Robert W. Cowan, Fredrik I. Thege, Vidhi Chandra, Sonja M. Wörmann, Hajar Rajaei, Prerna Malaney, Olivereen Le Roux, Sara L. Manning, Jack Hashem, Jennifer Bailey-Lundberg, Florencia McAllister, Andrew D. Rhim

## Abstract

Adenosine deaminase acting on RNA 1 (ADAR1) is an RNA-binding protein that deaminates adenosine(A) to inosine(I). A-to-I editing alters post-transcriptional RNA processing making ADAR1 a critical regulator of gene expression. Consequently, Adar1 has been implicated in organogenesis. To determine the role of Adar1 in pancreatic development and homeostasis, we specifically deleted *Adar1* from the murine pancreas (*Ptf1a^Cre/+^; Adar1^Fl/Fl^*). The resulting mice had stunted growth likely due to malabsorption associated with exocrine pancreas insufficiency. Analyses of pancreases revealed ductal expansion, heightened interferon-stimulated gene expression and an increased influx of immune cells. In addition, we observed an increased prevalence of CD4^+^ T and natural killer cells in their splenic tissue. These results indicate an association between loss of pancreatic Adar1 with dysregulation of systemic immunity. Concurrent deletion of *Adar1* and *Mavs*, a signaling protein implicated in the innate immune pathway rescued the degenerative phenotype and resulted in normal pancreatic development. Taken together, our work suggests that the primary function of Adar1 in the pancreas is to prevent aberrant activation of the Mavs-mediated innate immune pathway, thereby maintaining pancreatic homeostasis.

**Summary statement:** This work defines the role of Adar1 in pancreatic development and homeostasis.

## INTRODUCTION

Epigenetic alterations to RNA are widespread and can influence numerous cellular processes (Roundtree et al., 2017). RNA editing is an epigenetic modification wherein RNA nucleoside bases are modified by cellular enzymes. Adenosine-to-inosine (A-to-I) editing is the most prevalent form of RNA editing, in which A-to-I hydrolytic deamination is catalyzed by adenosine deaminase acting on RNA (ADAR) enzymes (Eisenberg and Levanon, 2018).

In mammals, there are three Adar proteins. Adar1 is expressed ubiquitously in all tissues (Kim et al., 1994) while Adar2 and Adar3 are primarily expressed in the brain (Melcher et al., 1996). *Adar1* encodes a constitutively expressed p110 isoform, localized in the nucleus, and an interferon-inducible p150 isoform that shuttles between the nucleus and cytoplasm (Patterson and Samuel, 1995).

Adar proteins have nonredundant functions, as demonstrated by the different phenotypes of *Adar1*^-/-^, *Adar2*^-/-^, and *Adar3*^-/-^ mice. Adar1 is essential for murine embryogenesis as *Adar1*^-/-^, *Adar1p150* ^-/-^, and editing-deficient *Adar1^E816A/E816A^* mice die *in utero* and display overexpression of interferon-stimulated genes (ISGs) (Wang et al., 2004, Hartner et al., 2004, Liddicoat et al., 2015, Ward et al., 2011). Moreover, these mice display widespread apoptosis, liver disintegration, and impaired hematopoiesis. In the absence of Adar1-mediated editing, melanoma differentiation-associated protein 5 (Mda5; encoded by *Ifih1*), a pattern recognition receptor, recognizes unedited endogenous RNA (Ahmad et al., 2018) and activates mitochondrial anti-viral signaling (Mavs) protein, its downstream effector. Mavs-related pathways include induction of ISGs and stimulation of innate immunity. Adar1 is a specific negative regulator of the Mda5-Mavs pathway and as such, *Adar1*^-/-^*Mavs*^-/-^ and *Adar1*^-/-^*Ifih1*^-/-^ mice do not exhibit exaggerated ISG expression and can be rescued to birth (Mannion et al., 2014, Pestal et al., 2015).

Adar1 exerts Mavs-dependent and independent functions as *Adar1*^-/-^Mavs^-/-^ mice exhibit developmental defects in the kidney and spleen among other organs (Pestal et al., 2015). Mavs-dependent functions are predicted to be RNA editing-mediated, whereas Mavs-independent functions are attributed to the regulation of microRNA biogenesis (Ota et al., 2013), RNA processing (Bahn et al., 2015), and protein recoding (Pestal et al., 2015). Organ-specific knockout models have revealed that Adar1 is essential for development and homeostasis of the liver, thymus, and hematopoietic compartment (Ben-Shoshan et al., 2017, Wang et al., 2015, Hartner et al., 2009, Vongpipatana et al., 2020); however, its role in the pancreas remains unknown.

The pancreatic exocrine compartment consists of highly specialized acinar cells that synthesize zymogens (amylase, lipase, trypsin, and chymotrypsin) necessary for digestion, and ductal cells that express carbonic anhydrase and secrete bicarbonate, which aids in neutralizing acidic chyme and pancreatic juice in the duodenum (Cano et al., 2007, Grapin-Botton, 2005). The islets of Langerhans comprise the pancreatic endocrine compartment, which produce hormones to regulate blood glucose levels (Pandiri, 2014).

We sought to determine Adar1’s role in pancreatic development and homeostasis by conditionally deleting *Adar1* from the pancreas.

## RESULTS and DISCUSSION

### Pancreas-specific *Adar1* deletion results in a runted phenotype and progressive exocrine pancreatic degeneration

Pancreas-associated transcription factor 1a (Ptf1a) is expressed on embryonic day 10.5 (E10.5) in multipotent pancreatic progenitors (Kawaguchi et al., 2002). To understand Adar1’s role in pancreatic development and homeostasis, we generated mice with the following genotype: Ptf1a^<u>C</u>re/+^(“C”); Rosa26^LSL-<u>Y</u>FP^(“Y”); <u>A</u>dar1^Fl/Fl^(“A”), herein denoted as CYA (**Figure 1A**). *Adar1* recombination was detected in the CYA pancreas (**Figure 1B**), but not in the spleen or kidney (**Figure S1A**), confirming the organ-specificity of the genetic model. At birth, control and CYA mice were indistinguishable in appearance; however, pancreas-specific *Adar1* deletion led to a runted phenotype evident at day 4 (D4) postpartum (**Figure S1B**), and very prominent at D21 (**Figure 1C**). Notably, CYA mice had significantly lower body mass compared to controls (**Figure 1D**). At E18.5 and at birth, the CYA mice were present at expected Mendelian ratios, which excluded the possibility of embryonic lethality (**Figure S1C**).

**Figure 1.**
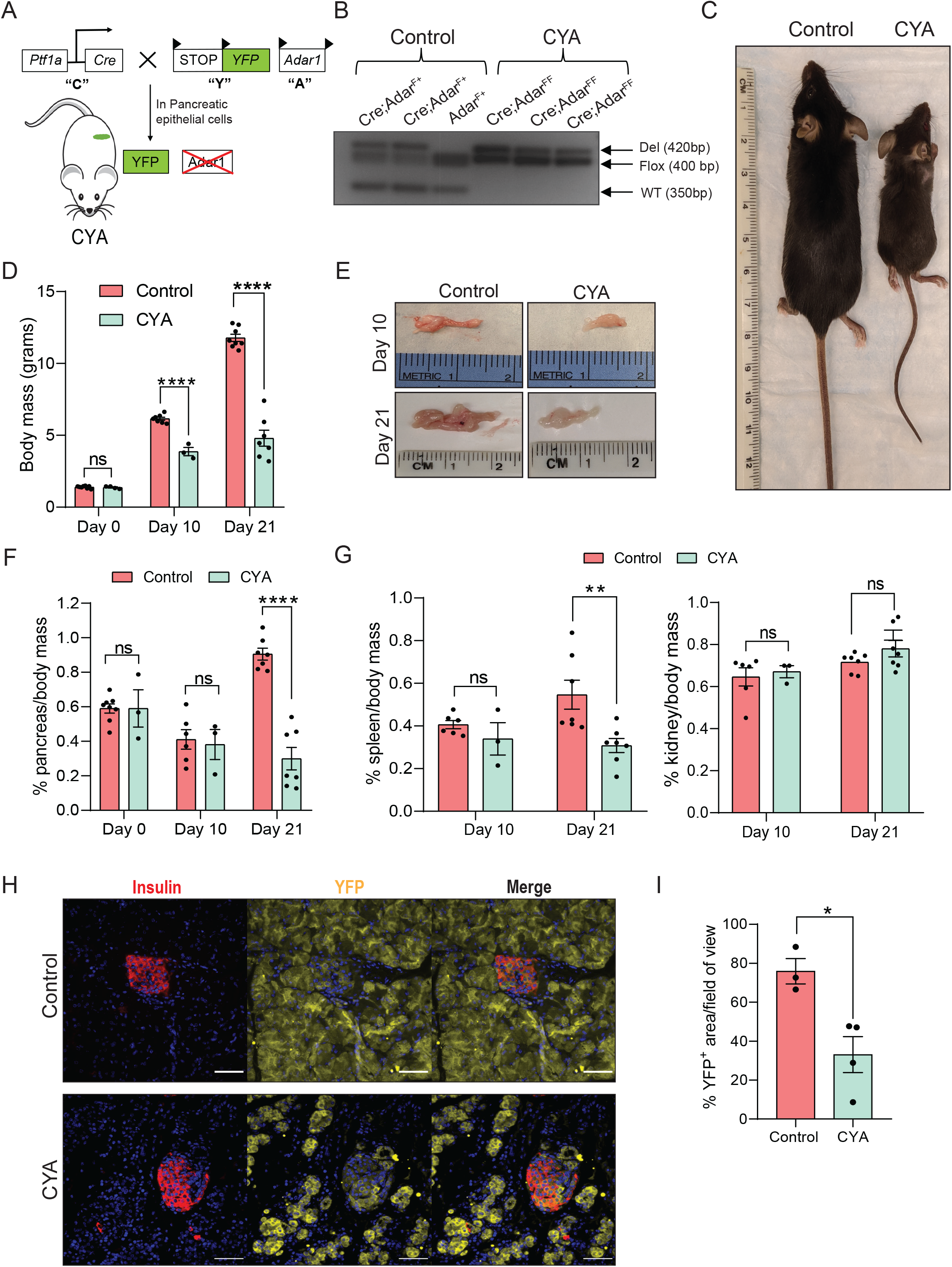
Conditional knockout of *Adar1* in the pancreas leads to a runted phenotype. **(A)** Schematic of the Ptf1a^<u>C</u>re/+^ Rosa26^LSL-<u>Y</u>FP^ <u>A</u>dar1^Fl/Fl^ (CYA) mouse model. **(B)** Control and CYA pancreases were assayed for *Adar1* recombination at birth. **(C)** Representative images of control and CYA mice. **(D)** Body mass of control and CYA mice at D0, D10 and D21. **(E)** Representative images of pancreases from control and CYA mice at D10 (top) and D21 (bottom). **(F)** Relative pancreatic mass (% pancreas/body mass) at D0, D10 and D21. **(G)** Relative spleen (left) and kidney (right) mass of control and CYA mice at D10 and D21. **(H)** Immunofluorescence exhibiting insulin and YFP in control and CYA pancreata at D21. Scale bar: 50 μM. **(I)** Quantification of YFP^+^ area in (H).

We next isolated and evaluated pancreatic tissue. Compared to controls, CYA mice had visibly smaller pancreases at D10 and D21, but not at D0 (**Figures 1E and S1D**). At D21, the relative pancreatic mass of CYA mice was significantly lower compared to controls **(Figure 1F)**. Likewise, at D21, the relative splenic mass, but not kidney mass, was reduced in CYA mice compared to controls **(Figure 1G).** We noted YFP^+^ cells in both the endocrine (insulin^+^) and exocrine compartment of the pancreas; however, the YFP^+^ area in the CYA pancreas was markedly reduced compared to controls (**Figure 1H and I**).

To understand if reduced pancreatic mass is a result of defective development or progressive degeneration, we assessed the control and CYA pancreases microscopically at E18.5, D2, D10, and D21. Histopathological analysis showed that the acinar cell area in CYA mice was comparable to that in controls at E18.5; however, it significantly reduced by D2 (**Figures 2A and B**). Additionally, CYA pancreases exhibited an increase in duct-like structures and an extensive immune infiltrate. Therefore, we stained for keratin19 (Ck19), a ductal cell marker, and observed an increase in the number of Ck19^+^ clusters in the CYA pancreases compared to controls at D10 (**Figures 2C and D**). We observed a similar amylase^+^ area (exocrine pancreas) in control and CYA pancreases at E18.5, however by D21 the exocrine area in CYA mice was drastically lower, confirming the progressive loss of the acinar cell area (**Figures 2E and F**). A higher number of apoptotic nuclei was observed in the CYA pancreases compared to controls at D21 (**Figures 2G and H**), which suggests that the decreased pancreatic mass in the CYA mice is attributable to apoptosis. Lastly, to quantify immune infiltrate observed via histology, we stained for CD45, a pan-leukocyte marker, and observed a significantly higher number of CD45^+^ cells in CYA pancreases compared to controls at D21 (**Figures 2I and J**). Our results suggest that the CYA pancreases develop normally; however, Adar1 is required for pancreatic acinar cell homeostasis and that the loss of Adar1 leads to progressive and rapid pancreatitis, and pancreatic degeneration due to apoptosis.

**Figure 2.**
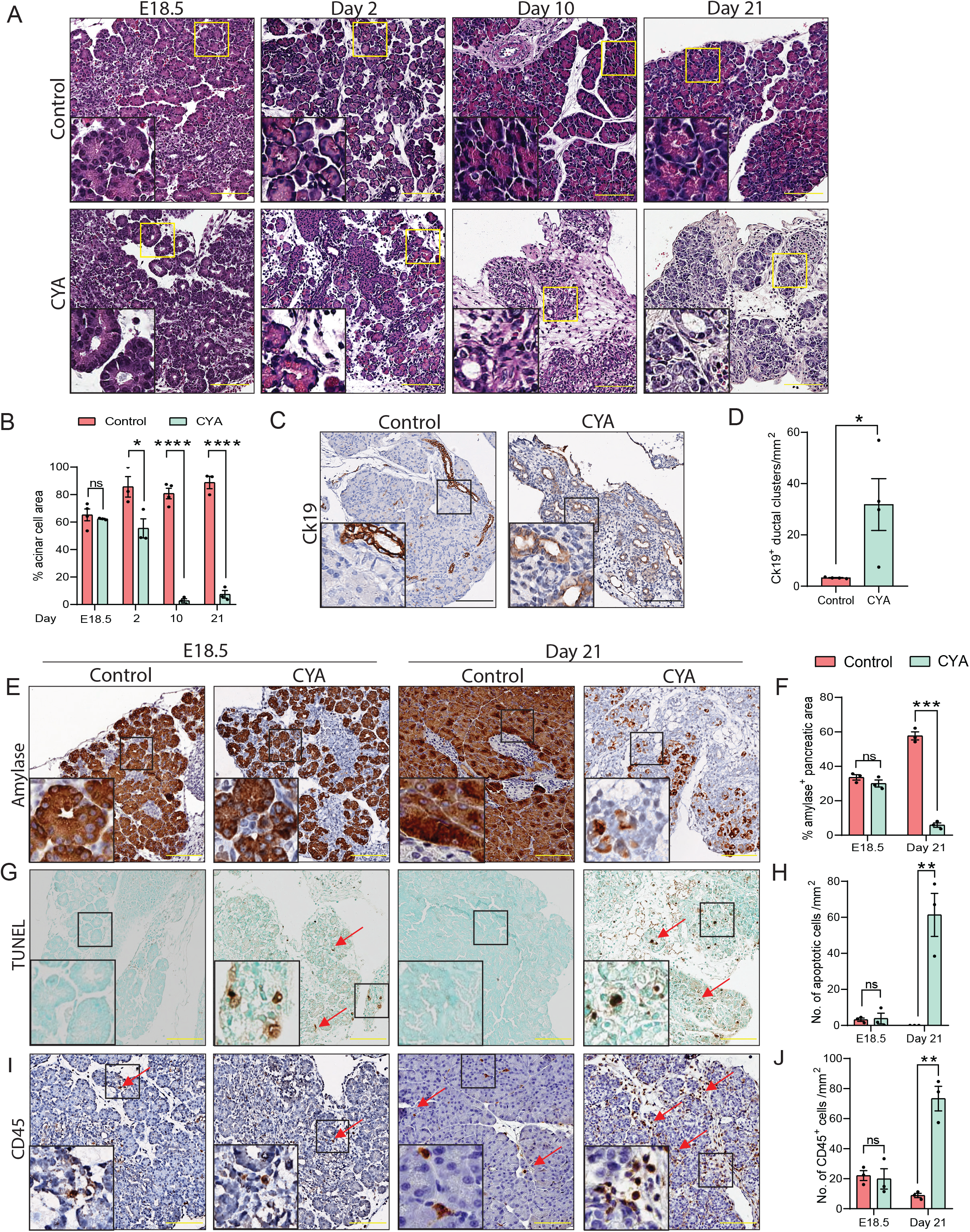
*Adar1* deletion results in loss of acinar cells and pancreatitis. **(A)** Histology of control and CYA pancreases at embryonic day 18.5 (E18.5), D2, D10 and D21. **(B)** Quantification of acinar cell area in (A). **(C)** Control and CYA pancreases at D10 assayed for keratin19 (Ck19) ^+^ clusters (red arrows) and **(D)** its quantification. **(E)** Representative images and **(F)** quantification of amylase^+^ area in control and CYA pancreata**. (G)** Representative images and **(H)** quantification of apoptotic nuclei (red arrows) in control and CYA pancreases**. (I)** Representative images and **(J)** quantification of CD45^+^ cells (red arrows) in control and CYA pancreata. Scale bar: 100 μM.

### Loss of Adar1 results in exocrine pancreatic insufficiency and malabsorption

Since the pancreas is required for digestion of food and blood glucose homeostasis, we hypothesized that pancreatic degeneration results in dysfunction contributing to the runted phenotype of CYA mice. Thus, we assessed whether loss of Adar1 affects the pancreatic exocrine and endocrine compartments equally. Surprisingly, we observed a relative increase in the insulin^+^ area (endocrine compartment) of the CYA pancreases compared to controls (**Figure S2A and B**), very likely due to their lower total pancreatic area. In our model, loss of Adar1 specifically affects the pancreatic exocrine but not endocrine compartment, which is not unusual, as it has been previously reported that cells arising from common progenitors have a differential response to *Adar1* deletion (Gacem et al., 2020, Liddicoat et al., 2016). However, we analyzed the pancreas at D21 and effects of *Adar1* deletion on the endocrine compartment may be apparent at later time points. Role of Adar1 in the pancreatic endocrine compartment can be further studied using an endocrine-specific Cre (Magnuson and Osipovich, 2013).

Pancreatic injury results in alteration of serum pancreatic enzymes and hormone concentrations (Lee and Papachristou, 2019, Galicia-Garcia et al., 2020), thus we next assessed their concentration in the mice. CYA mice exhibited significant reduction in serum lipase concentration but similar amylase, glucagon and insulin concentrations (**Figure S2C-F**) compared to controls at D21. We predicted that decreased lipase concentration would hinder digestion of complex triglycerides and fat absorption from the intestine resulting in higher-than-normal stool fat content, known as steatorrhea. Indeed, fecal fat content from CYA stool samples had a significantly higher number of unabsorbed fat globules than control samples, in which unabsorbed fat was absent (**Figure S2G-H**). Our results suggest that loss of Adar1 causes pancreatic exocrine insufficiency, leading to malabsorption and steatorrhea, symptoms reminiscent of patients suffering from pancreatic insufficiency (Lindkvist, 2013). The malabsorption of nutrients suggests that CYA mice are malnourished and provides an explanation for their runted phenotype

### Pancreas-specific *Adar1* knockout induces systemic immune dysregulation

The CYA spleens were significantly smaller (splenic hypoplasia) compared to those of controls (**Figure 1G**), thus we evaluated whether this was due to increased apoptosis but found no difference in the number of apoptotic nuclei (**Figures S3A and B**). The spleen can initiate immune responses in recognition of host damage or infection (Lewis et al., 2019); therefore, we assessed whether the CYA spleens had an aberrant immune cell composition. We observed a significantly higher proportion of CD4^+^ T cells and NK cells in CYA spleens compared to controls (**Figures S3C, D and S4**). Loss of Adar1 is associated with type 1 interferon signaling (Wang et al., 2015, Hartner et al., 2009), which modulates T and NK cell responses (McNab et al., 2015, Brinkmann et al., 1993, Madera et al., 2016). Aberrant numbers of CD4^+^ T cells and NK cells are implicated in numerous autoimmune disorders (Palmer and Weaver, 2010, Skapenko et al., 2005, Liu et al., 2021). Due to the anatomical proximity of the pancreas and spleen, certain chemokines and cytokines released from the Adar1 knockout pancreas could be leading to splenic hypoplasia and changes in the lymphocyte population, though they remain to be identified. Our findings suggest that loss of Adar1 in pancreas results in local inflammation that triggers systemic immune activation.

### Concurrent deletion of *Mavs* prevents the pancreatic degeneration caused by the loss of Adar1

*Adar1* deletion stimulates a Mavs-mediated innate immune pathway and induces expression of ISGs in *Adar1*^-/-^ mice (Mannion et al., 2014, Pestal et al., 2015). We found a higher expression of ISGs in CYA pancreases compared to controls at D2 (**Figure 3A**), consistent with activation of the Mavs pathway. Adar1 has a Mavs-dependent role in innate immune activation, and its Mavs-independent functions can influence murine organ development (Vongpipatana et al., 2020, Pestal et al., 2015). Therefore, we hypothesized that co-deletion of *Mavs* would prevent innate immune activation and suppress aberrant ISG expression, but that pancreas may still have developmental defects as described in other organs. Accordingly, we generated mice with double knockout of *Adar1* and *Mavs* from the pancreas (denoted as CYAM): Ptf1a^<u>C</u>re/+^(“C”); Rosa26^LSL-<u>Y</u>FP^(“Y”); <u>A</u>dar1^Fl/Fl^ (“A”); <u>M</u>avs^-/-^(“M”). As expected, ISG expression in CYAM mice was similar to controls, except *Mx1* which was significantly lower (**Figure 3B**). Surprisingly, CYAM mice developed similarly to the control mice. Macroscopically, concurrent deletion of *Mavs* rescued the runted phenotype of CYA mice, and CYAM body mass did not differ from controls at 6 or 12 months (**Figures 3C, 3D and S5A**). We monitored CYAM mice for 12 months for delayed developmental defects, but none were noted. Furthermore, there was no difference in the relative pancreatic mass of CYAM and control mice at 6 or 12 months (**Figures 3E, 3F, and S5B**). To exclude the possibility that CYAM pancreases developed normally because they expressed Adar1, we confirmed Adar1 recombination in the pancreas at 2 months (**Figure 3G**). Microscopic examination of the CYAM pancreases showed normal architecture devoid of apoptotic cells or excess immune infiltrate (**Figures. 3H-L**). Our work suggests that concurrent *Mavs* knockout completely rescues the degenerative phenotype of the CYA mice, and that CYAM pancreases develop normally. However, we cannot exclude the possibility of transcriptomic differences or changes in protein translation in CYAM mice (Bajad et al., 2020).

**Figure 3.**
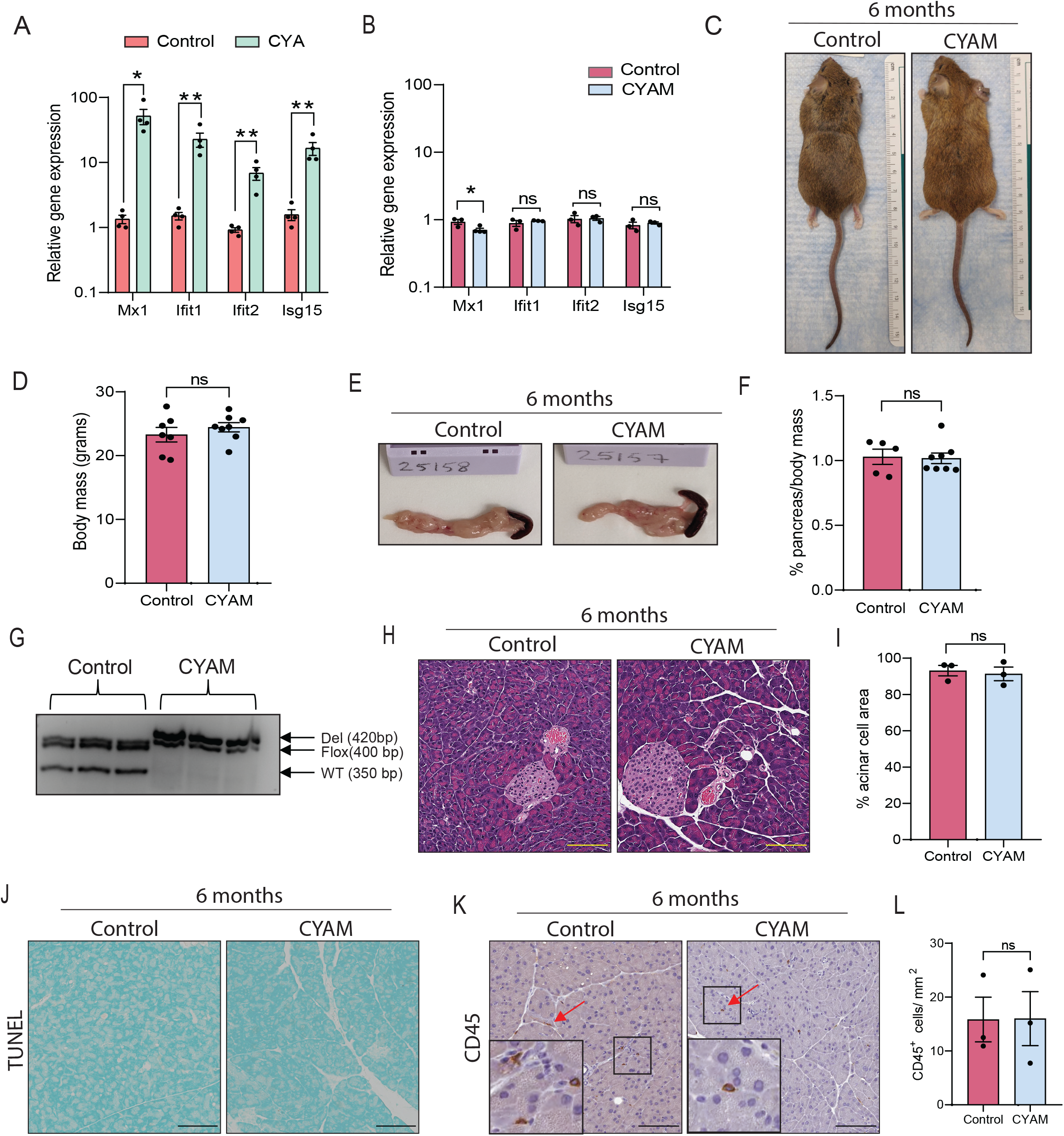
Concurrent *Mavs* knockout can rescue the runted phenotype of the CYA mice. **(A)** Quantitative PCR (qPCR) for a panel of interferon-stimulated genes (ISGs) from control and CYA pancreata at D2. **(B)** qPCR for ISGs from control and CYAM pancreata at 2 months. **(C)** Representative images and **(D)** quantification of body mass of control and CYAM mice at 6 months. **(E)** Representative images and **(F)** quantification of relative pancreatic mass of control and CYAM mice at 6 months. **(G)** Pancreases from control and CYAM mice were assayed for *Adar1* recombination. **(H)** Representative images of control and CYAM pancreases and **(I)** quantification of acinar cell area. **(J)** Representative images of apoptotic nuclei and **(K, L)** + CD45^+^ (red arrows) cells in control and CYAM pancreases. Scale bar: 100 μM.

Additionally, we generated mice heterozygous for Mavs, denoted as CYAM^**+/-**^. The body mass of CYAM^+/-^ mice was comparable to that of control and CYAM mice at 12 months (**Figure S5A**), however, their relative pancreatic mass was significantly lower (**Figure S5B**). Microscopic examination showed a significantly reduced acinar cell area in the CYAM^+/-^ pancreases compared to controls (**Figure S5C and D**). Our results suggest that concurrent deletion of *Mavs* can rescue the degenerative phenotype observed in CYA mice, but presence of even one *Mavs* allele can sub-optimally trigger innate immunity pathways leading to slower pancreatic degeneration in the absence of Adar1.

### Co-deletion of *Mavs* restores pancreatic function that is disrupted by Adar1 deletion

Since we determined that the CYAM pancreases were similar to those of controls in appearance and histology, we next evaluated whether CYAM mice had any deficiencies arising from pancreatic dysfunction. Therefore, we assessed whether the pancreatic exocrine and endocrine compartments had developed normally in CYAM mice. Immunohistochemistry analysis showed similar exocrine and endocrine area in the CYAM mice compared to controls (**Figure 4A-D**). We then evaluated serum amylase, lipase and glucagon concentrations. Contrary to results from CYA mice, serum lipase concentrations were comparable between CYAM mice and controls (**Figure 4E-G**), suggesting that *Mavs* deletion rescued the pancreatic dysfunction of CYA mice. Additionally, we tested pancreatic function by conducting oral glucose tolerance tests (Andrikopoulos et al., 2008) and assessing fat absorption from the intestine (Drummey et al., 1961). When challenged with glucose, both groups were equally adept at insulin secretion and maintenance of glucose homeostasis **(Figure 4H and I)**. In contrast to steatorrhea observed in CYA mice, we did not observe any unabsorbed fat globules in the stools of the CYAM mice **(Figure 4J)**. Using these criteria, our results suggest that CYAM pancreases have no functional deficiencies.

**Figure 4.**
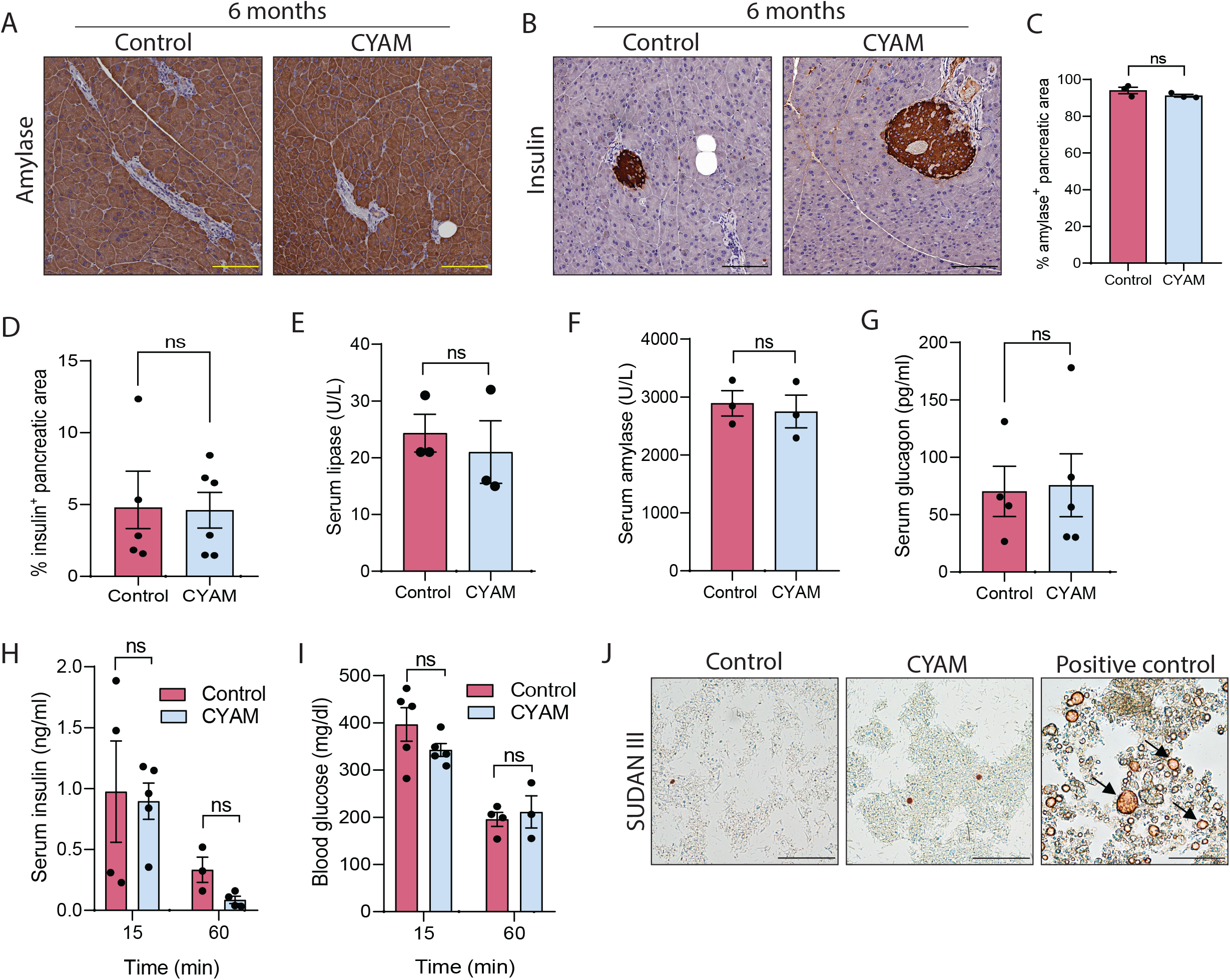
Co-deletion of *Mavs* can restore pancreatic function. Representative images of **(A)** amylase^+^ and **(B)** insulin^+^ area and **(C, D)** their respective quantification. Scale bar: 100 μM. **(E)** Serum lipase, **(F)** amylase, and **(G)** glucagon concentration in control and CYAM mice. **(H)** Serum insulin and **(I)** blood glucose concentration, 15 and 60 mins post-oral glucose administration. **(J)** Representative images of control and CYAM fecal fat assayed for fat globules with SUDAN III. Scale bar: 50 μM.

To our knowledge this is the first study that reports the pancreatic function of Adar1. The absence of Adar1 in pancreatic cells leads to induction of ISGs and pancreatitis, similar to type I interferonopathies observed in patients suffering from Aicardi-Goutieres syndrome, a subset of which harbor *ADAR1* genetic mutations (Crow and Manel, 2015). Our results are also in accordance with previous studies, which have reported increased apoptosis and ISG expression in *Adar1*^-/-^ mice and in organ-specific *Adar1* knockout models (Ben-Shoshan et al., 2017, Hartner et al., 2004, Hartner et al., 2009, Wang et al., 2004). Counter to Adar1’s previously reported role in the development of thymus, kidney and spleen (Pestal et al., 2015, Vongpipatana et al., 2020), our investigation suggests that loss of Adar1 does not affect pancreatic development. Our results indicate that Adar1 has a unique function in different cell lineages and that its functions in one organ cannot be extrapolated to another organ.

In summary, our work shows that the Mavs-mediated function of Adar1 is essential for homeostasis and the suppression of interferon signaling in the pancreas.

## MATERIALS AND METHODS

### Mouse models

A pancreas-specific *Adar1* knockout model was generated by crossing the *Adar1* floxed allele (*<u>A</u>dar1^Fl^*; “A”) (Mutant Mouse Resource & Research Centers stock no: 34619-JAX) with the pancreas-specific Ptf1a^<u>C</u>re/+^(“C”) model (Kawaguchi et al., 2002). Genotyping and *Adar1* recombination assessment were performed as described previously (Wang et al., 2004). The *Rosa26^LSL-<u>Y</u>FP^* (“Y”) reporter allele (The Jackson Laboratory, Stock No: 006148) was introduced to enable lineage tracing (Srinivas et al., 2001, Rhim et al., 2012). *<u>M</u>avs*^-/-^ (“M”) mice (Sun et al., 2006) were obtained from The Jackson Laboratory (stock no: 008634) and genotyped according to the protocols available on their website. For a pancreas-specific Adar1 knockout, we generated Ptf1a^<u>C</u>re/+^(“C”); Rosa26^LSL-<u>Y</u>FP^(“Y”); <u>A</u>dar1^Fl/Fl^ (“A”) mice, herein denoted as CYA. For a double knockout of *Adar1* and *Mavs*, we generated Ptf1a ^<u>C</u>re/+^(“C”); Rosa26^LSL-<u>Y</u>FP^(“Y”); <u>A</u>dar1^Fl/Fl^(“A”); <u>M</u>avs^-/-^ (“M”) mice, herein denoted as CYAM. The controls used are either CYA^F+^, CY, YA^FF^, CYM or M^-/-^. All animal experiments were conducted in compliance with the National Institutes of Health guidelines for animal research and were approved by the Institutional Animal Care and Use Committee of The University of Texas MD Anderson Cancer Center.

### Histopathology

Mice were euthanized, their tissues were isolated and weighed. For relative organ mass (% organ mass/body mass) was calculated for each mouse. Tissues were then fixed in zinc-formalin and embedded in paraffin, and 5-μm sections were obtained and stained with hematoxylin (Dako) and eosin (VWR) following standard protocols.

### Immunohistochemistry

Formalin-fixed, paraffin-embedded tissue sections were deparaffinized in xylene and rehydrated in graded ethanol. Antigen retrieval was performed in a citrate buffer (pH 6.0). Slides were treated with 3% hydrogen peroxide for 15 min and blocked for an hour with either CAS-block (ThermoFisher; #8120) or 2.5% goat serum. Tissue sections were incubated overnight at 4°C with primary antibodies against mouse amylase, insulin, keratin19 and CD45 (antibody details, including dilutions, can be found in **Table S1)**. A SignalStain DAB Substrate Kit (Cell Signaling Technology) was used for chromogenic antibody detection. Slides were counterstained with hematoxylin and mounted in Acrymount (StatLab). Apoptotic nuclei were detected according to the manufacturer’s instructions using an *in situ* apoptosis detection kit (Abcam, #ab206386) which utilizes principals of the terminal deoxynucleotidyl transferase nick-end labeling (TUNEL) assay.

Slides were imaged at 20X magnification on the Aperio CS2 Scanner and analyzed with the Aperio eSlide Manager software.

Positive cells/staining area were quantified and normalized to the total pancreatic area per field of view. A minimum of 5 different fields were assessed, and the average number of positive cells/area for each biological replicate was reported.

### Immunofluorescence

Formalin-fixed, paraffin-embedded tissue sections were deparaffinized in xylene and rehydrated in graded ethanol. Antigen retrieval was performed in a citrate buffer (pH 6.0). Sections were then blocked with blocking buffer (10% normal goat or donkey serum, 1% bovine serum albumin, 0.4% Triton X-100 in phosphate-buffered saline) for an hour and incubated overnight at 4°C with primary antibodies against green fluorescent protein (cross-reactive with yellow fluorescent protein) and insulin (**Table S1**). Nuclei were counterstained with 4’,6-diamidino-2-phenylindole (DAPI). TrueVIEW Autofluorescence Quenching Kit (Vector Laboratories, #SP-8400) was used according to the manufacturer’s instructions to quench autofluorescence. Epifluorescent images were obtained at a 40X magnification on an Olympus IX73 inverted fluorescent microscope.

### Serum enzyme and blood glucose detection

Serum amylase and lipase concentrations were determined at the Clinical Pathology core facility at MD Anderson Cancer Center with an Integra 400 Plus (Roche Diagnostics). Serum insulin and glucagon concentrations were measured according to the manufacturer’s instructions with an Ultra-Sensitive Mouse Insulin ELISA Kit (Crystal Chem, #90080) and a Mouse Glucagon ELISA Kit (Crystal Chem, #81518).

For fasting blood glucose, mice were fasted for 4 hours and blood glucose was measured using a Roche Diagnostics Accu-Chek Inform II glucometer. Oral glucose tolerance test was performed as described previously (Andrikopoulos et al., 2008). Briefly, mice fasted overnight (16h) and were given an oral bolus of 2 mg/kg of glucose. Retro-orbital blood was collected 15 min and 1h after glucose administration. Blood glucose and serum insulin were measured as described above.

### Fecal fat quantification

Protocol for fecal fat assessment was adapted from a previously described protocol (Drummey et al., 1961). Briefly, 25mg of stool was added to a 1:1 solution of 95% ethanol and deionized water, and vortexed. 25μL of supernatant was mixed with 10μL of a saturated solution of SUDAN III (Sigma-Aldrich, # S4131-25G) in 95% ethanol. Fat globules were quantified (under 40X magnification) as the number of fat globules per field of view. A minimum of five different fields were assessed, and the average number of fat globules per biological replicate was reported.

### Flow cytometry

Single cell splenocytes were isolated by passing the spleen through a 40μm mesh. Cells were treated with Fixative-Free Lysing Solution, High-Yield Lyse (Invitrogen, #HYL250) for RBC lysis. Samples were then stained with a Live/Dead Blue dye, followed by incubation with Fc Block (BD Biosciences) and then stained with anti-mouse Nk1.1, Ly6G, Ly6C, CD19, CD11b, CD11c, CD3, CD8, CD4, and CD45 antibodies (**Table S1)**. Sample acquisition was performed on an LSRFortessa X-20 Cell Analyzer flow cytometer (BD Biosciences). The analysis was performed with FlowJo software, version 10 (FlowJo LLC). Results are expressed as relative percentage of total gated live CD45^+^ cells.

### RNA isolation and quantitative polymerase chain reaction (qPCR)

RNA was isolated using the Qiagen RNeasy Mini Kit and reverse transcribed using the SuperScript III First-Strand Synthesis System (Invitrogen, #18080-51). qPCR was performed using SYBR Green PCR Master Mix (Lifetech, #43-676-59) on a ViiA 7 Real-Time PCR System (Applied Biosystems). The primers used are listed in **Table S2**. Beta-2-microglobulin was used for normalization. Assays were performed with 3 technical replicates.

### Statistical analysis

All statistical analyses were performed using Prism software (GraphPad Software, Inc.). Data is expressed as the mean plus or minus the standard error of the mean (SEM). Statistical significance between two groups was assessed using a two-sample test, unless otherwise noted. p<0.05 was considered statistically significant. Ns = not significant, *=p<0.05, ***=p < 0.001 ** = p < 0.01, **** = p < 0.0001.

## ACKNOWLEDGEMENTS

We thank members of the Rhim and McAllister lab for critical reading of the manuscript, as well as Laura L. Russell, scientific editor, Research Medical Library, for helpful editorial review. We also thank the technical staff at the Clinical Pathology core facility and the Flow Cytometry and Cellular Imaging Core Facility at MD Anderson Cancer Center for their help and expertise. This shared resource is partially funded by NCI Cancer Center Support Grant P30CA16672.

